# Cytokine control of systemic hypoxia tolerance in *Drosophila*

**DOI:** 10.1101/2025.06.30.661614

**Authors:** Kate Ding, Prajakta Bodkhe, Byoungchun Lee, Danielle Polan, Amy Wycislik, Tiffany Cheung, Sophie Wu, Savraj S Grewal

## Abstract

Systemic hypoxia—reduction in oxygen supply to all tissues and organs—occurs in both physiological and pathological conditions in animals. Under these conditions, organisms must adapt their physiology to ensure proper functioning and survival. While extensive research has characterized how individual cells sense and adapt to low oxygen conditions, the mechanisms that coordinate whole-body responses to systemic hypoxia remain poorly understood. In this study, we uncover an interorgan signaling response mediated by the cytokine Unpaired-3 (upd3) that is important for systemic hypoxia tolerance in *Drosophila*. We demonstrate that hypoxia rapidly induces upd3 expression and activates JAK/STAT signaling in both larvae and adults. Interestingly, we discovered a sex-specific requirement for this pathway, with females requiring upd3 for hypoxia survival while males do not. We also identify the intestine as a critical source of hypoxia-induced upd3 and show that gut-derived upd3 signals to the fat body and oenocytes to mediate hypoxia tolerance by promoting nitric oxide synthase expression. Furthermore, we reveal an unexpected role for the canonical hypoxia response transcription factor HIF-1α/sima as a molecular brake, preventing lethal upd3 overproduction, revealing that hypoxia survival requires precise cytokine dosage control. Our findings define a gut-to-fat/oenocyte signaling axis that coordinates systemic hypoxia adaptation, highlighting the complex interplay between classic hypoxia response pathways and cytokine signaling in maintaining organismal homeostasis during oxygen limitation. This work provides important insights into how organisms adapt to systemic hypoxia, with potential implications for understanding systemic hypoxia-related human pathologies such as respiratory disorders and sleep apnea.

## Introduction

Oxygen is essential for aerobic life, serving as the terminal electron acceptor in cellular respiration that powers energy production across the animal kingdom. When oxygen availability becomes limited—a condition known as hypoxia—organisms must rapidly adapt their physiology and metabolism to maintain homeostasis (Semenza, 2011). These adaptations are critical not only during normal animal development and in specific ecological niches but also in numerous pathological conditions that result in oxygen deprivation such as stroke and lung disorders (Bickler and Buck, 2007; Jahani et al., 2020; Ramirez et al., 2007; Samanta et al., 2017; Semenza, 2014b; Stupnikov and Cardoso, 2017).

Although atmospheric air contains ∼20% oxygen, cells and tissues typically experience significantly lower oxygen levels under physiological conditions. For example, the early fetus develops in an environment where oxygen levels can be as low as 1% (Dunwoodie, 2009), while oxygen levels in adult tissues can vary from 1 to 10% depending on the organ (McKeown, 2014). In many diseases, oxygen supply becomes further restricted, either locally (as in tumors, ischemia, and stroke) or systemically (as in respiratory disorders, COVID-19, and sleep apnea) (May and Mehra, 2014; Semenza, 2011; 2014a; b; Xie and Simon, 2017). Systemic hypoxia results from insufficient oxygen reaching all of the body’s tissues and organs, potentially leading to multi-organ failure and life-threatening complications. Extensive work in cell culture has established how individual cells sense and respond to low oxygen, particularly how they modulate gene expression and reprogram metabolism to survive hypoxic conditions (Holdsworth and Gibbs, 2020; Nakazawa et al., 2016; Schito and Rey, 2018; Xie and Simon, 2017). In contrast, responses to systemic hypoxia in whole animals represents a fundamentally different challenge that requires systems-level coordination (Baik and Jain, 2020; Midha et al., 2023; Samanta *et al*., 2017). These responses likely rely on inter-organ communication networks to integrate oxygen sensing across multiple organs, but the nature of these networks and how they operate remain largely unexplored.

*Drosophila melanogaster* provides an excellent model system for investigating organismal adaptive responses to hypoxia (Callier et al., 2015; Callier et al., 2013; Farzin et al., 2014; Harrison et al., 2018; Harrison and Haddad, 2011). In their natural ecology, *Drosophila* larvae grow in rotting, fermenting food rich in microorganisms—an environment characterized by low ambient oxygen (Callier *et al*., 2015; Markow, 2015). They have therefore evolved multiple sophisticated mechanisms to tolerate hypoxia. One key pathway involves the conserved hypoxia-sensing transcription factor HIF-1α (known as sima in *Drosophila*), which is stabilized under low oxygen conditions and subsequently activates genes involved in metabolic adaptation and oxygen delivery (Centanin et al., 2008; Centanin et al., 2005; Gorr et al., 2006; Lavista-Llanos et al., 2002; Romero et al., 2007; Texada et al., 2019). *Drosophila* research has also pioneered our understanding of inter-organ communication networks, revealing how specific tissues can detect environmental changes and signal to other organs to coordinate whole-body physiological responses (Alvarez-Ochoa et al., 2021; Boulan et al., 2015; Kannangara et al., 2021; Kim et al., 2021b; Leopold and Perrimon, 2007; Malita and Rewitz, 2021; Medina et al., 2022; Meschi and Delanoue, 2021; Okamoto and Watanabe, 2022; Owusu-Ansah and Perrimon, 2015; Rajan and Perrimon, 2011; Texada et al., 2020; Yoon et al., 2023). Thus, while all cells can adjust their metabolism in response to changes within their local environment, these tissue-level sensors ensure that physiological adaptations to environmental conditions are coordinated across multiple organs according to the organism’s overall needs. This has been best exemplified in the context of how flies adapt to fluctuations in dietary nutrient availability. Extensive work has shown how tissues such as the gut, fat body, and brain couple sensing of nutrients to behavioral, physiological, and metabolic adaptations through a complex inter-organ signaling network (Droujinine and Perrimon, 2016; Koyama et al., 2020; Medina *et al*., 2022; Meschi and Delanoue, 2021; Tennessen and Thummel, 2011). A few reports have indicated that similar tissue-tissue signaling responses may also underlie organismal adaptation to hypoxia. For example, sima activation in the larval fat body triggers fat-to-brain signals that suppress insulin-like peptide production, reducing growth under oxygen-limited conditions (Texada *et al*., 2019), while sima induction in specialized neurons modifies neuroendocrine signaling to stimulate blood cell production in hypoxia (Cho et al., 2018). Nevertheless, the inter-organ networks coordinating systemic hypoxia adaptation remain poorly understood.

The Unpaired (upd) family of cytokines are key regulators of interorgan communication in *Drosophila* (Zandawala and Gera, 2024). These secreted signaling molecules activate the conserved JAK/STAT pathway by binding to their cell surface receptor, Domeless (Agaisse and Perrimon, 2004). Upds play an important role in mediating communication between tissues to maintain metabolic and physiological homeostasis, especially in response to fluctuations in nutrient availability (Ingaramo et al., 2020; Rajan et al., 2017; Rajan and Perrimon, 2012; Zhao and Karpac, 2017). Additionally, upds can be induced by a variety of stressors including pathogenic infection, oxidative stress, nutrient stress, and tissue damage (Hersperger et al., 2024; Srinivasan et al., 2016; Yang et al., 2015). Importantly, the source of upd production varies with the specific stress context, as does the target tissue and the resulting physiological effects. This context-dependent signaling allows upds to orchestrate complex, multi-tissue responses that coordinate activities across the gut, fat body, brain, cardiac tissue, and immune cells to control tissue repair, metabolism, endocrine signaling, and growth (Brent and Rajan, 2020; Cai et al., 2021; Chakrabarti et al., 2016; Ding et al., 2021; Gera et al., 2022; Hersperger *et al*., 2024; Houtz et al., 2017; Huang et al., 2020; Ingaramo *et al*., 2020; Jiang et al., 2009; Kierdorf et al., 2020; Liu et al., 2025; Liu et al., 2024; Nagai et al., 2023; Obata et al., 2018; Osman et al., 2012; Rai et al., 2025; Rajan *et al*., 2017; Rajan and Perrimon, 2012; Romao et al., 2021; Shin et al., 2020; Sodders et al., 2025; Srinivasan *et al*., 2016; von Frieling et al., 2020; Wang et al., 2014; Woodcock et al., 2015; Yang *et al*., 2015; Zhao and Karpac, 2017; Zhou et al., 2013).

A critical aspect of upd signaling is its context-dependent role in stress responses, which can be either adaptive or detrimental to organismal fitness. In acute stress conditions, upd induction often promotes beneficial protective mechanisms. For instance, transient activation of the upds in gut epithelial cells following injury or infection stimulates stem cell-mediated tissue regeneration (Buchon et al., 2009; Jiang *et al*., 2009; Obata *et al*., 2018; Osman *et al*., 2012; von Frieling *et al*., 2020; Zhou *et al*., 2013). Similarly, injury-induced upd3 from hemocytes supports tissue repair and survival (Chakrabarti *et al*., 2016). Additionally, in normal physiology, upd3 production from pericardial cells maintains ECM protein expression essential for cardiac function (Gera *et al*., 2022). However, these same signaling pathways become harmful when chronically activated or in pathological contexts. For example, persistent upd activation in the aging intestine leads to epithelial dysplasia and shortened lifespan (Li et al., 2016). Upon high-fat diet consumption hemocyte-derived upd3 disrupts glucose homeostasis and reduces longevity (Woodcock *et al*., 2015), while fat body-derived upd2 causes nephrocyte dysfunction (Zhao et al., 2025). Additionally, age-related upd3 upregulation in oenocytes contributes to cardiac dysfunction (Huang *et al*., 2020). Perhaps most strikingly, in tumor models, upd3 secretion triggers systemic pathologies including blood-brain barrier compromise and metabolic dysregulation that accelerate mortality (Ding *et al*., 2021; Kim et al., 2021a; Liu *et al*., 2025). This dual nature—beneficial in physiological contexts but potentially harmful when dysregulated— underscores the importance of investigating upd signaling in specific physiological and pathological conditions to understand its precise roles in stress adaptation.

Here, we investigate how upd3 signaling mediates whole-body hypoxia tolerance in *Drosophila*, revealing an interorgan communication network that coordinates systemic hypoxia adaptation through the interplay of cytokine and classic hypoxia response pathways.

## RESULTS

### Hypoxia induces upd3/JAK/STAT signaling

We first examined whether hypoxia induces the cytokine/JAK/STAT signaling pathway. To do this, we subjected *w^1118^* adult flies and larvae to different durations of either hypoxia (1% oxygen) or normoxia (ambient air) and then collected samples for gene expression analysis by qRT-PCR. We found that hypoxia exposure led to a strong upregulation of *upd3* mRNA levels in both male and female adult flies (Figure 1A, B), an effect that was apparent within 4hrs of hypoxia exposure. To explore whether this upregulation led to functional induction of cytokine signaling we also measured the mRNA levels of well-described STAT target genes as a functional readout of JAK/STAT pathway activation. This analysis revealed strong upregulation of the STAT target gene *SOCS36E* in males and females with a similar timecourse (Figure 1C, D). We also saw that another class of STAT regulated genes, the *Turandot* genes, *TotA, TotC, TotM,* were also induced upon hypoxia exposure in adults (Figure S1A, B). Previous studies have shown that flies become immobilized and cease feeding during exposure to 1% oxygen. However, nutrient starvation for a 16-hour period failed to induce *upd3* mRNA in both male (Figure S1C) and female (Figure S1D) animals suggesting that cessation of feeding is not responsible for *upd3* induction in hypoxia. Finally, we examined whether the hypoxia induction of upd/JAK/STAT signaling also occurred in larvae. We exposed larvae to 2, 4, and 8 hours of normoxia or hypoxia and collected them for gene expression analysis. Similar to our observations in adults, we detected induction of both *upd3* and *SOCS36E* mRNA in hypoxic larvae (Figure 1E, F). Together, these results indicate that induction of upd3 cytokine signaling is a robust response to hypoxia exposure in *Drosophila*.

**Figure 1:**
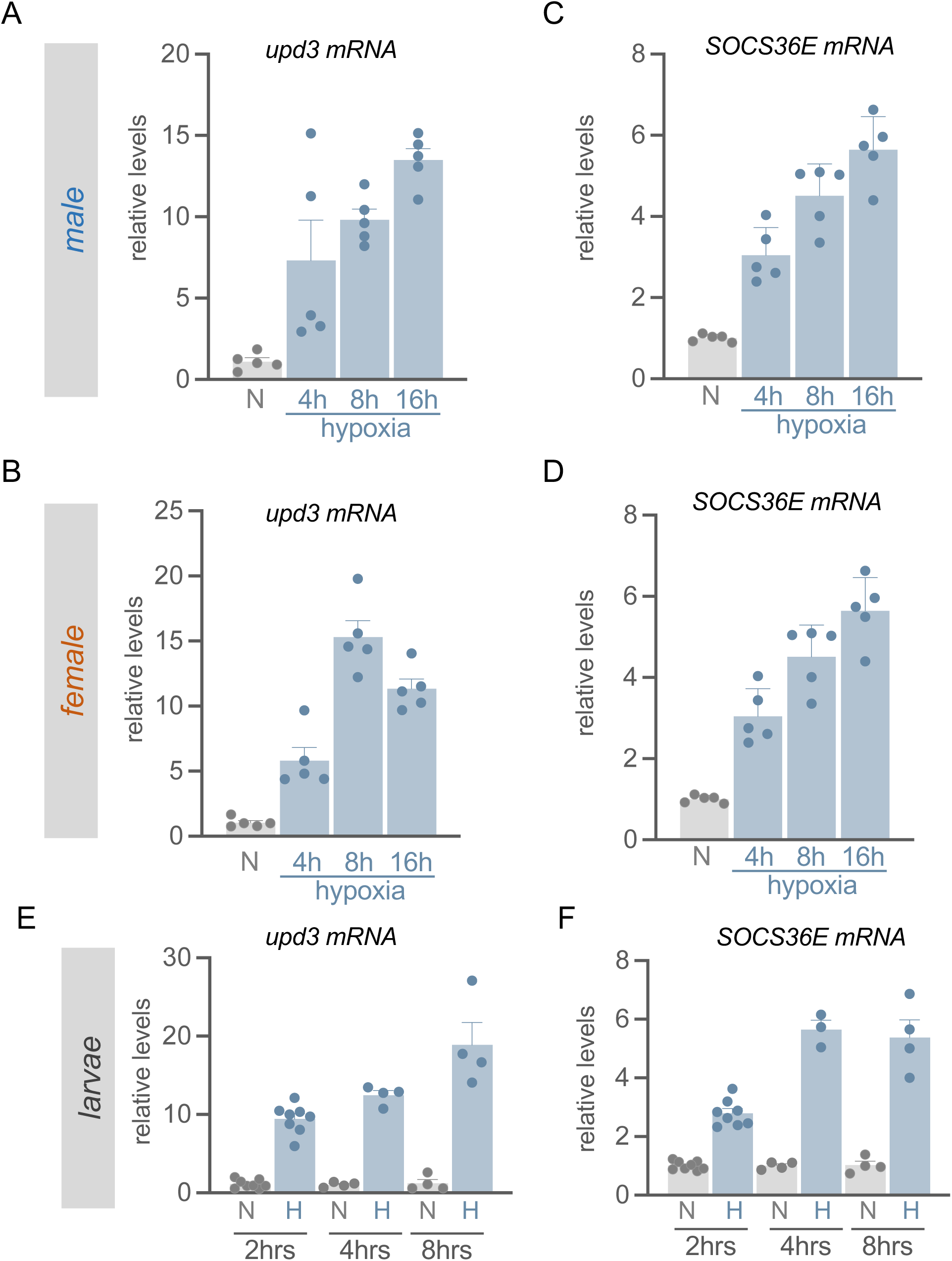
Hypoxia induces upd3/Jak/STAT signaling in larvae and adults. (A-D) Age-matched flies (*w^1118^*) were maintained in normoxia or hypoxia (1% oxygen), then collected for gene expression analysis. qRT-PCR measurement of *upd3* in A) males and B) females following 4, 8, and 16 hours of hypoxia. STAT-target gene *SOCS36E* measured in C) males and D) females were measured following 4, 8 and 16 hours of hypoxia. Data represents mean + SEM, N=5 independent groups of samples (5 animals per group) per experimental condition. Data points represent independent samples. E, F) qRT-PCR measurements of E) *upd3* mRNA and F) *SOCS36E* mRNA in larvae exposed to 2, 4 and 8 hours of hypoxia (1% oxygen). Data represents mean + SEM, N=3-8 independent groups of animals (10 animals per group) per experimental condition. Data points represent independent samples.

### Sex-specific requirement for upd3 in hypoxia tolerance

We next investigated the functional relevance of increased upd3 signaling in hypoxia tolerance. To test this, we examined survival under hypoxic conditions in a *upd3*-null mutant line (*upd3Δ)* (Osman *et al*., 2012). We exposed age-matched 7-day-old control (*w^1118^*) and *upd3* mutant adult flies to an acute bout of hypoxia that resulted in 50-80% survival of control flies. Our survival analysis revealed that female *upd3*-null mutants exhibited significantly reduced survival compared to their control counterparts (Figure 2B). In contrast, *upd3*-null mutant males showed no decrease in hypoxia survival compared to control males and even displayed a small but significant increase in survival (Figure 2A). Given that female metabolism is altered by mating status (Ahmed et al., 2020; Hadjieconomou et al., 2020; Hudry et al., 2016; Reiff et al., 2015; Zipper et al., 2020), we asked whether mating status affected the requirement for *upd3* in hypoxia tolerance. Both mated and virgin female *upd3*-null animals showed decreased survival in hypoxia compared to controls (Figure S2B, C).

**Figure 2:**
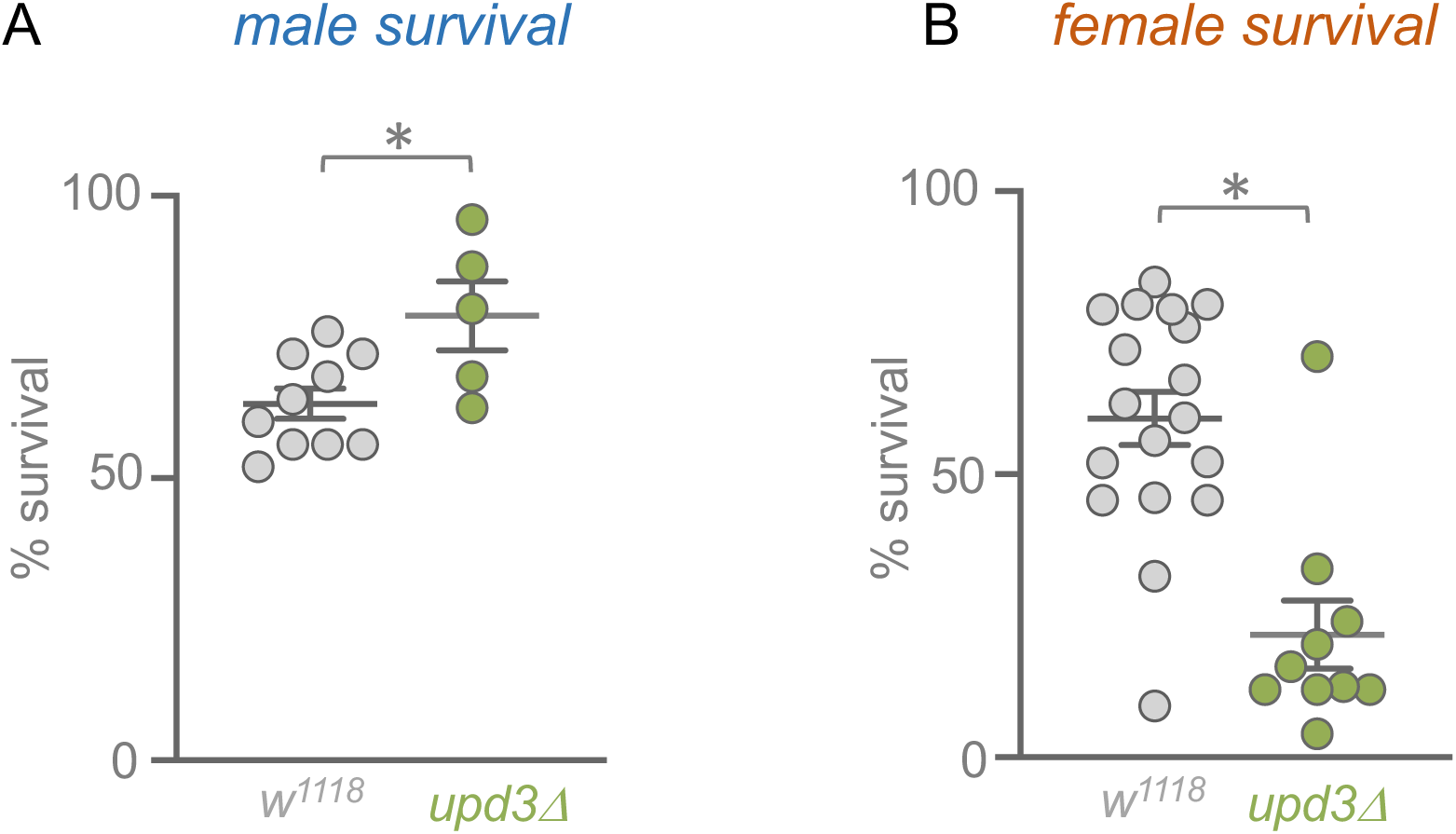
upd3 is required for hypoxia tolerance in females but not males. Hypoxia survival measured in control (*w^1118^*) vs upd3-null (*upd3Δ*) A) males and B) females. Data represents mean ± SEM, N≥5 independent groups of animals (15-25 per group) per experimental condition. *p < 0.05, unpaired *t*-test. Data points represent independent samples.

These results suggest a sexually dimorphic requirement for *upd3* in mediating hypoxia tolerance. While *upd3* is necessary for hypoxia tolerance in females regardless of mating status, it appears dispensable or even detrimental in males. Further studies will be needed to determine if this sex difference occurs through the sex determination pathway, which is emerging as an important mediator of fly physiology. However, for the remainder of this study, we focused on exploring how upd3 mediates hypoxia tolerance in mated females.

### Intestinal upd3 coordinates systemic hypoxia responses

The upds facilitate tissue-to-tissue communication to control adaptive responses to stress. They are expressed in different tissues and can act in an autocrine or paracrine manner. Therefore, we wanted to investigate what key tissue(s) produce upd3 to promote hypoxia tolerance. One key tissue known to produce upd3 in response to stress is the adult midgut. Previous studies have shown how a variety of different stresses including infection, tissue damage and nutrient stress can upregulate intestinal upd3 to mediate both local and systemic adaptive response (Buchon *et al*., 2009; Cai *et al*., 2021; Houtz *et al*., 2017; Jiang *et al*., 2009; Nagai *et al*., 2023; Obata *et al*., 2018; Takeishi et al., 2013; von Frieling *et al*., 2020; Zhou *et al*., 2013).

When we exposed control (*w^1118^*) females to 16 hours of normoxia or hypoxia and dissected guts for qRT-PCR analysis, we found a strong upregulation of *upd3* mRNA levels in hypoxia (Figure 3A). We also analyzed flies carrying an *upd3* GFP transcriptional reporter (*upd3Gal4, UAS-GFP*) that were exposed to either normoxia or hypoxia. Microscopic visualization of the intestines from these flies showed that hypoxia-exposed animals had stronger GFP expression in their guts compared to normoxia controls, which was especially evident in the R2 region of the intestine (Figure 3B). Moreover, this increased GFP expression was mostly seen in the large epithelial enterocytes. To interrogate this finding further, we knocked down upd3 using an enterocyte-specific driver (*mex>upd3-RNAi*) and measured intestinal *upd3* mRNA levels using qPCR. We found that enterocyte upd3 knockdown resulted in significant reduction (∼80%) of *upd3* mRNA (Figure 3C), suggesting that the enterocytes are the main intestinal cell type upregulating upd3 in hypoxia. Furthermore, when we repeated this experiment but now measured whole-body *upd3* mRNA levels, we saw that enterocyte upd3 knockdown led to a significant ∼40% decrease in the hypoxia-mediated upregulation of whole-body *upd3* levels (Figure 3D). These results suggest that, although upd3 is likely upregulated in several tissues, the intestinal enterocytes contribute a large proportion of the systemic upregulation of upd3 in hypoxia-exposed flies.

**Figure 3:**
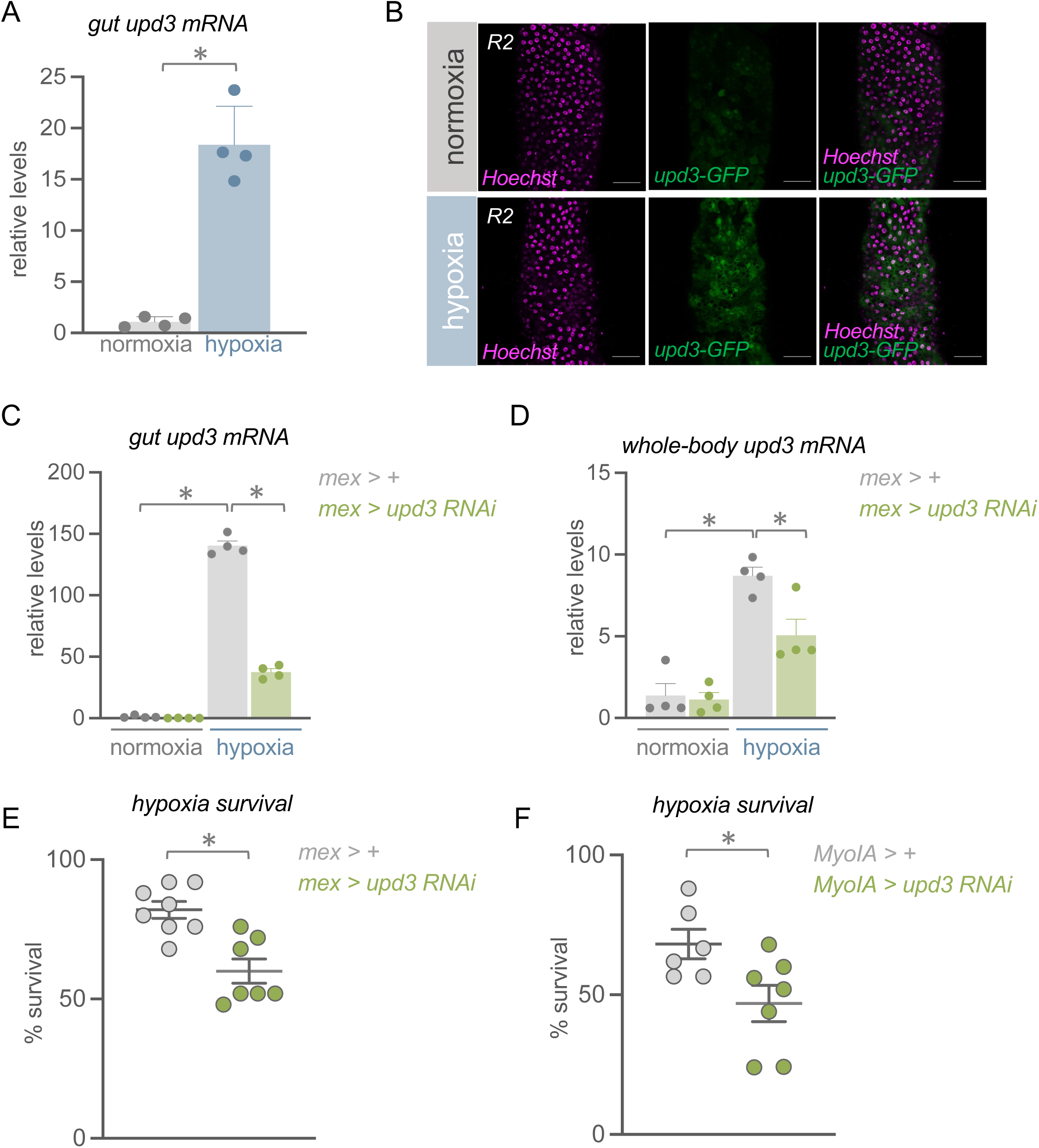
Gut-derived upd3 promotes hypoxia tolerance. A) qRT-PCR analysis of *upd3* mRNA of guts from *w^1118^* adult females maintained in normoxia or exposed to 16 hours of hypoxia (1% oxygen). RNA was isolated from whole guts. Data represents mean + SEM, N=4 independent groups of samples (5 intestines per group) per experimental condition. Data points represent independent samples. B) Female flies expressing GFP under control of an upd3 promoter (*upd3-gal4, UAS-GFP*) were exposed to hypoxia (1% oxygen) or maintained in normoxia for 16 hours. Whole guts were dissected, stained with Hoechst to visualize nuclei, and mounted for visualization of GFP expression. Representative images of R2 region shown. Scale bar = 50µm. C) qRT-PCR measurement of *upd3* mRNA in whole intestines with control (*mex > +*) vs enterocyte upd3 knockdown flies (*mex > upd3-RNAi)* in hypoxia. Data represents mean + SEM, N=4 independent groups of samples (5 intestines per group) per experimental condition. *p < 0.05, Student’s *t*-test following 2-way ANOVA. Data points represent independent samples D) *upd3* mRNA also measured in whole-body. Data represents mean + SEM, N=4 independent groups of samples (5 animals per group) per experimental condition. *p < 0.05, Student’s *t*-test following 2-way ANOVA. Data points represent independent samples. E, F) Hypoxia survival measured in enterocyte *upd3* knockdown animals using two different drivers (*mex > upd3-RNAi, Myo1A > upd3-RNAi*) compared to their respective control genotypes (*mex > +, Myo1A > +*). Data represents mean ± SEM, N≥6 independent groups of animals (15-25 per group) per experimental condition. *p < 0.05, unpaired *t*-test. Data points represent individual samples.

We next investigated the functional role for gut-derived upd3 in hypoxia tolerance. We subjected control flies (*mex>+*) and flies with gut-specific *upd3* knockdown (*mex>upd3-RNAi*) to hypoxia and measured their survival. We saw that suppressing intestinal *upd3* expression significantly reduced fly survival compared to controls (Figure 3E). We repeated this experiment using another enterocyte-specific driver, *Myo1A-Gal4*, to knock down *upd3* and observed similar results with the *Myo1A>upd3-RNAi* flies showing significantly reduced survival compared to controls (*Myo1A>+*) (Figure 3F). To rule out potential developmental effects of upd3 knockdown, we employed a temperature-sensitive enterocyte driver to induce RNAi knockdown of *upd3* specifically during adulthood. These adult-specific upd3 knockdown flies (*Myo1A^ts^>upd3-RNAi*) also showed reduced survival under hypoxic conditions compared to controls (*Myo1A^ts^>+*) (Figure S2C). Collectively, these results demonstrate that the gut serves as an important source of hypoxia-induced *upd3* expression essential for hypoxia tolerance.

### Gut-derived upd3 signals to fat and oenocytes to promote hypoxia tolerance

Having established the intestine as an important source of expression for upd3 in mediating hypoxia tolerance, we next investigated which tissue(s) gut-derived upd3 might be communicating with to stimulate JAK/STAT signaling and mediate this hypoxia tolerance response. We performed qRT-PCR on tissue from isolated heads, thoraces, abdomens (with guts removed), and ovaries from female *w^1118^* flies exposed to either normoxia or hypoxia to examine expression of the STAT target gene, *SOCS36E*. We saw that *SOCS36E* was upregulated in all tissues in hypoxia-exposed flies suggesting widespread stimulation of JAK/STAT signaling in low oxygen. However, the hypoxia induction of *SOCS36E* was most pronounced in isolated abdominal tissues (Figure 4A). We also saw strong induction of two other STAT targets genes, *TotA* and *TotM,* in abdominal tissues of hypoxia-exposed flies (Figure S3A). We then examined whether gut-derived upd3 can signal to abdominal tissues, by inducing acute *upd3* expression from intestinal enterocytes (*mex^ts^>UAS-upd3*) and measuring expression of STAT target genes in isolated abdominal tissues. Compared to controls (*mex^ts^>+*), we observed induction of both *TotA* and *TotM* mRNA in the abdomens of *mex^ts^>UAS-upd3* (Figure 4B) indicating that gut-derived upd3 can stimulate JAK/STAT signaling in abdominal tissues.

**Figure 4:**
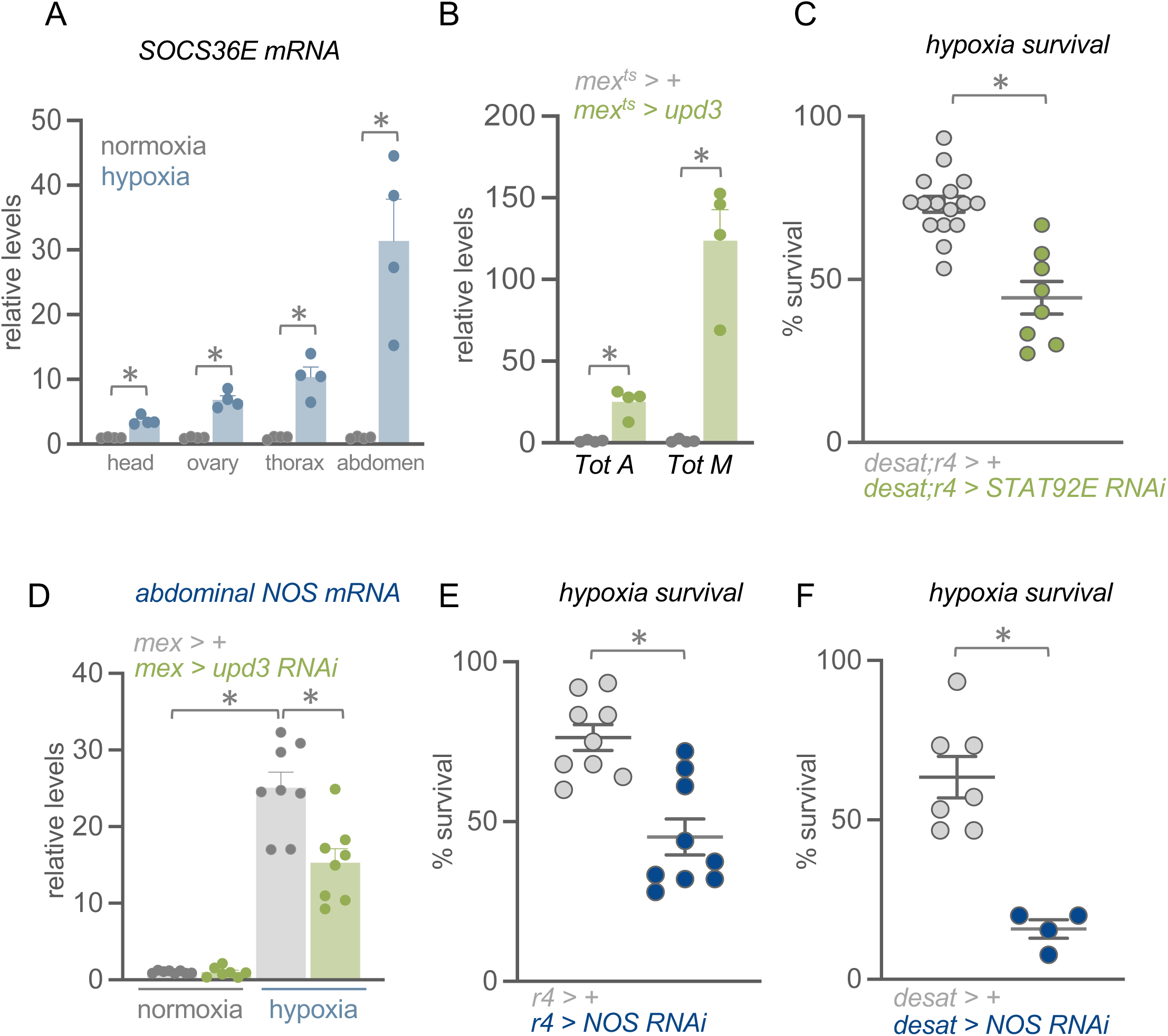
Gut-derived upd3 controls adipose expression of hypoxia regulators. A) Adult females (*w^1118^*) were maintained in normoxia or exposed to 16 hours of hypoxia. Different tissues including the head, ovary, thorax and abdomen were dissected for *SOCS36E* mRNA expression analysis. Data represents mean + SEM, N=4 independent groups of samples (5-10 tissues per group) per experimental condition. *p < 0.05, unpaired *t*-test. Data points represent independent samples. B) qRT-PCR analysis of STAT target genes (*Tot A* and *Tot M)* in the adult female abdominal tissue with intestinal upd3 overexpression (*mex^ts^ > upd3*) vs control (*mex^ts^ > +*). Data represents mean + SEM, N=4 independent groups of samples (5 abdomens with intestines and ovaries removed per group) per experimental condition. *p < 0.05, ns not significant, Student’s *t*-test following 2-way ANOVA. Data points represent independent samples. C) Survival measured in animals with RNAi knockdown of STAT92E in the two main abdominal tissue types – oenocytes and fat body – (*desat;r4 > STAT92E-RNAi*) vs control (*desat;r4 > +*). Data represents mean ± SEM, N≥8 independent groups of animals (15-25 per group) per experimental condition. *p < 0.05, unpaired *t*-test. Data points represent individual samples. D) *NOS* mRNA levels measured in abdomens when *upd3* was knocked down from intestinal enterocytes (*mex > upd3-RNAi*) vs control (*mex > +*). N=4 independent groups of samples (5 abdomens with intestines and ovaries removed per group) per experimental condition. *p < 0.05, Student’s *t*-test following 2-way ANOVA. Data points represent independent samples. Hypoxia survival measured with NOS knockdown in E) fat body alone (*r4 > NOS RNAi*) vs control (*r4 > +*), or F) oenocytes alone (*desat > NOS RNAi*) vs control (*desat > +*). Data represents mean ± SEM, N≥4 independent groups of animals (15-25 per group) per experimental condition. *p < 0.05, unpaired *t*-test. Data points represent individual samples.

Adult abdominal tissue is composed mostly of fat body and oenocytes (Ghosh et al., 2020). These tissues are functionally equivalent to mammalian adipose and liver and play important roles in lipid and sugar storage, mobilization and metabolism, as well as functioning as endocrine tissues (Arrese and Soulages, 2010; Bland, 2022; Ghosh *et al*., 2020; Huang et al., 2022; Kuhnlein, 2011; Martinez et al., 2020; Meschi and Delanoue, 2021; Sriskanthadevan-Pirahas et al., 2022; Stefana et al., 2017; Yamada et al., 2018). Through these effects both the fat body and oenocytes have well-established roles in coordinating whole-body physiological responses to changes in environmental stimuli such as food, pathogens and toxins. In particular, previous studies have shown that the fat body is a key target of endocrine upd3 signaling involved in the regulation of systemic physiology and stress responses (Gera *et al*., 2022; Liu *et al*., 2025). We therefore assessed the functional importance of abdominal STAT signaling in hypoxia tolerance. We generated a line carrying two independent gal4 drivers, *desat-Gal4* and *r4-Gal4* that drive expression specifically in oenocytes and fat body respectively. We used this line to express dsRNA against STAT92E (*desat;r4>STAT92E-RNAi*) simultaneously in both oenocytes and fat body and measured hypoxia survival. We found that oenocyte/fat body-specific STAT92E knockdown significantly reduced survival under hypoxic conditions compared to control animals (Figure 4C). Together these results suggest that one way that gut-derived upd3 can promote hypoxia tolerance is through signaling to the fat body and oenocytes.

### Gut-derived upd3 controls abdominal expression of nitric oxide synthase, a regulator of hypoxia survival

Nitric oxide (NO), which is synthesized by the enzyme nitric oxide synthase (Nos), is a conserved mediator of hypoxic responses and in *Drosophila* has been shown to be required for several aspects of hypoxia adaptation including cell cycle arrest, altered behaviour and survival (DiGregorio et al., 2001; Dijkers and O’Farrell, 2009; Teodoro and O’Farrell, 2003; Wingrove and O’Farrell, 1999). Interestingly, we found that *Nos* mRNA expression levels were strongly elevated in isolated abdominal tissues from hypoxia-exposed flies suggesting a potential hypoxic role for Nos in these tissues. We therefore examined whether Nos might be a downstream effector of gut-to-oenocyte/fat body upd3 signaling. We knocked down upd3 specifically in intestinal enterocytes (*mex>upd3-RNAi*) and then analyzed *Nos* expression in abdominal tissues from normoxic versus hypoxic flies. We saw that gut-specific upd3 knockdown significantly reduced the hypoxia-induced increase in abdominal *Nos* expression (Figure 4D). This regulation appeared to be tissue-specific, as we did not observe gut *upd3*-dependent regulation of *Nos* expression in other tissues including the head, thorax, and ovaries (Figure S3A-C). To determine whether abdominal Nos was functionally important for hypoxia tolerance, we used RNAi to knock down *Nos* in either the fat body or oenocytes. In both cases, *Nos* knockdown significantly reduced survival under hypoxic conditions compared to controls (Figure 4E, F). These results demonstrate that gut-derived upd3 promotes hypoxia tolerance by regulating *Nos* expression specifically in abdominal tissues, identifying Nos as a key downstream effector of the gut-to-fat/oenocyte signaling axis.

### HIF-1α limits upd3 signaling to promote hypoxia tolerance

We next explored how hypoxia regulates upd3 expression. Hypoxia-inducible factor 1 (HIF-1) is one of the best-characterized transcriptional regulators of hypoxia response and in mammalian cells has been shown to be pro-inflammatory in part through expression of cytokines (Nizet and Johnson, 2009; Palazon et al., 2014; Palsson-McDermott et al., 2015; Tannahill et al., 2013). We therefore investigated whether HIF-1α (known as sima in *Drosophila*) was required for upd3 induction during hypoxia. We used RNAi to knock down sima ubiquitously using the *da-gal4* driver (*da>sima-RNAi*) and compared *upd3* mRNA levels under hypoxic conditions to controls (*da>+*). Interestingly, we found that sima knockdown augmented the hypoxia-induced increase in *upd3* mRNA levels (Figure 5A) and led to a strong amplification of the hypoxia-induced increased in expression of the STAT target gene,*TotM* (Figure 5B). These results suggest that, rather than being required for upd3 induction, sima is needed to limit upd3/JAK/STAT signaling during hypoxia. This finding aligns with a prevailing theme in immunology: the necessity of fine-tuning cytokine signaling. While effective immune responses rely on rapid induction of cytokine signaling, it is equally important to limit excess cytokine signaling to avoid ‘cytokine-storm’ responses, whereby unchecked cytokine signaling can cause unwanted tissue damage and immunopathology (Cron et al., 2023; Fajgenbaum and June, 2020; Jahani *et al*., 2020; Medzhitov, 2021; Meizlish et al., 2021; Ye and Medzhitov, 2019). Based on our results, we hypothesized that one role of sima during hypoxia might be to limit the potentially deleterious effects of excessive upd3 levels to ensure proper adaptation to hypoxia. This hypothesis made two testable predictions: first, that overexpression of upd3 to create excess cytokine signaling would be detrimental in hypoxia; and second, that if sima’s role is to dampen excessive upd3 signaling, then lowering upd3 levels might partially protect against the lethality seen with sima knockdown. We tested both predictions.

**Figure 5:**
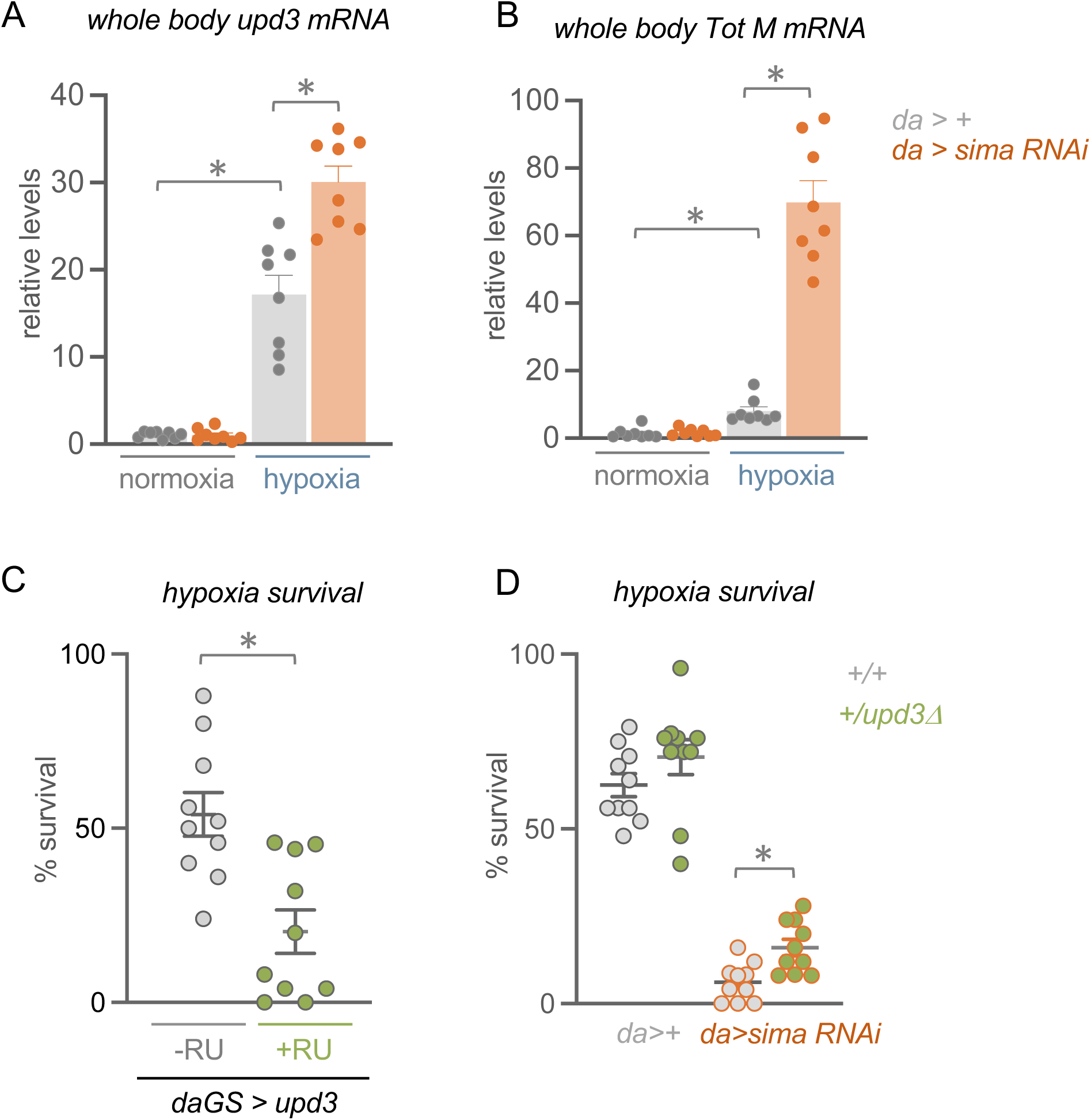
HIF-1 alpha/sima limits excess upd3/Jak/STAT signaling. Control (*da > +*) or sima knockdown (*da > sima-RNAi*) females were maintained in normoxia or exposed to hypoxia (1% oxygen) for 16 hours. Whole flies were then collected for qRT-PCR analysis. Whole body A) *upd3* mRNA and B) *TotM* mRNA were measured. Data represents mean + SEM, N=8 independent groups of samples (5 animals per group) per experimental condition. *p < 0.05, ns not significant, Student’s *t*-test following 2-way ANOVA. Data points represent independent samples. C) Hypoxia survival of control (*da-GS > upd3*, -RU486) vs ubiquitous *upd3* OE (*da-GS > upd3*, +RU486) adult flies. Data represents mean ± SEM, N=10 independent groups of animals (15-25 per group) per experimental condition. *p < 0.05, unpaired *t*-test. Data points represent individual samples. D) Control *(da > +)*, *upd3Δ* heterozygote (*da > upd3Δ*/+), *sima* knockdown (*da > sima-RNAi*) and *upd3Δ* heterozygote *sima* knockdown (*da > upd3Δ*/+*;sima-RNAi*) adult females were maintained in normoxia or exposed to hypoxia (1% oxygen) and survival measured. Data represents mean ± SEM, N≥8 independent groups of animals (15-25 per group) per experimental condition. *p < 0.05, unpaired *t*-test. Data points represent individual samples.

To test whether excess upd3 is detrimental in low oxygen, we used the GeneSwitch-Gal4 system to acutely overexpress *upd3* ubiquitously in adult female flies. *Upd3*-overexpressing flies (*daGS>upd3*, fed RU486) showed significantly reduced hypoxia survival compared to control flies (*daGS>upd3*, fed vehicle control) (Figure 5C). We confirmed this result using another ubiquitous inducible driver (*actGS-Gal4*) and observed the same outcome—excess *upd3* reduced hypoxia tolerance (Figure S4A)

We next tested whether reducing *upd3* gene dosage could rescue the lethality of sima knockdown flies. To lower *upd3* signaling, we carried out experiments in flies heterozygous for *upd3*. Using qRT-PCR, we first confirmed that the augmented *upd3* levels seen in hypoxic sima knockdown flies were reduced back to control hypoxic levels in *upd3* heterozygotes, confirming the effectiveness of reducing gene dosage of *upd3* in dampening the excess *upd3* expression (Figure S4B). We then tested hypoxia survival and found that sima knockdown flies heterozygous for the *upd3* deletion (*upd3Δ*/+*; da> sima-RNAi*) showed a small but significant improvement in survival under hypoxic conditions compared to sima knockdown flies with normal *upd3* levels (*da>sima-RNAi*) (Figure 5D). These results demonstrate that part of the lethality observed in *sima*-deficient flies is attributable to exaggerated *upd3* induction during hypoxia, supporting our hypothesis that sima functions to prevent harmful levels of cytokine signaling during hypoxia.

### Adipose HIF-1α regulates gut upd3 expression

Our experiments on sima focused on whole-body regulation of upd3, so we next turned to how sima might regulate intestinal upd3 production specifically. We repeated the experiment with ubiquitous sima knockdown but measured gut *upd3* mRNA levels directly. As with whole-body upd3, we observed augmented upd3 production in the gut under hypoxic conditions (Figure 6A), suggesting that sima’s modulation of whole-body upd3 levels may largely reflect augmentation of upd3 in the gut.

**Figure 6:**
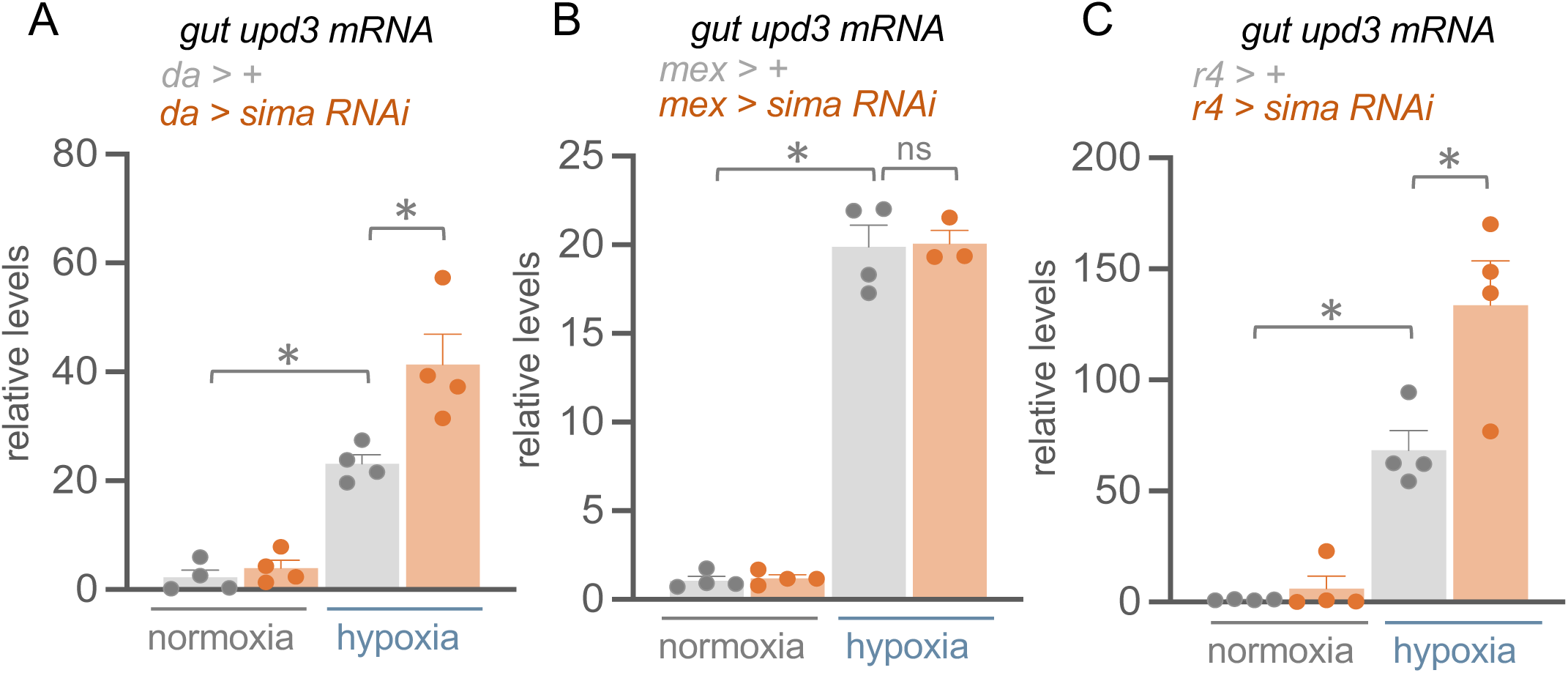
Adipose HIF-1 alpha/sima limits intestinal upd3 expression. Adult female flies of control and sima knockdown in A) the whole fly (*da > +, da > sima-RNAi*), B) gut (*mex > +, mex > sima-RNAi*) or C) fat body (*r4 > +, r4 > sima-RNAi*) were maintained in normoxia or hypoxia (1% oxygen) for 16 hours. Whole guts were immediately dissected for qRT-PCR analysis. Data represents mean + SEM, N=4 independent groups of samples (5 intestines per group) per experimental condition. *p < 0.05, ns not significant, Student’s *t*-test following 2-way ANOVA. Data points represent independent samples.

We initially hypothesized that sima in the gut might directly antagonize gut upd3 expression. To test this hypothesis, we generated enterocyte-specific sima knockdown flies and isolated their intestines after exposure to normoxia or hypoxia to measure *upd3* mRNA levels. Contrary to our expectation, gut-specific sima knockdown (*mex>sima-RNAi*) did not significantly alter intestinal *upd3* mRNA levels compared to controls (*mex>+*) under hypoxic conditions (Figure 6C). This finding suggested that gut sima does not directly regulate gut upd3 expression and that the effect of sima on upd3 is non-autonomous.

Previous work has shown that sima activity in the fat body during hypoxia plays an important non-autonomous role in controlling whole-body physiology, with effects thought to rely on sima modulation of fat-dependent endocrine signaling (Texada *et al*., 2019). The *Drosophila* fat body also serves as a central integration hub for signals to and from other tissues. Based on these observations, we examined the fat body as a potential source of sima-dependent regulation of intestinal upd3. We generated fat body-specific *sima* knockdown flies (*r4>sima-RNAi*), exposed them to hypoxia, and measured gut upd3 mRNA levels. Strikingly, intestines from fat body *sima* knockdown flies showed significantly greater induction of upd3 under hypoxic conditions compared to control (*r4>+*) intestines (Figure 6C). These results suggest that sima in the fat body acts as a hypoxia sensor that regulates upd3 expression in the gut, implying the existence of a sima-dependent signal mediating this tissue-to-tissue communication

We investigated whether reactive oxygen species (ROS) might mediate this interorgan communication. Since ROS are well-established inducers of upd3, we tested whether sima knockdown affects ROS-induced upd3 by exposing fat body-specific sima knockdown flies (*r4>sima-RNAi*) to paraquat, a ROS-generating chemical. We observed no significant difference in intestinal upd3 expression compared to controls (Figure S4C). Additionally, overexpressing the antioxidant enzyme Catalase A in the gut (*mex>CatA*) did not affect hypoxia-induced intestinal upd3 levels (Figure S4D). These results suggest that the hypoxia- and sima-dependent modulation of upd3 is independent of ROS signaling.

## DISCUSSION

Organisms must coordinate whole-body responses to environmental stresses through inter-organ communication networks. Our results reveal how cytokine signaling mediates such coordination during systemic hypoxia, identifying a gut-to-fat body axis that is essential for organismal survival under low oxygen conditions.

Upd3 has been shown to mediate adaptive responses to numerous environmental stresses including nutrient stress, infection, and toxins, and our work extends its role to the regulation of adaptation to low oxygen. We demonstrate that gut-derived upd3 is needed for survival in low oxygen, and our results suggest this occurs in part through signaling to two main tissues in the fly abdomen—the oenocytes and the fat body. Both cell types play important roles in energy metabolism, particularly in coordinating whole-body metabolic and physiological adaptations to changes in environmental conditions through their adipose- and liver-like roles in fat and sugar storage and mobilization, as well as their endocrine functions in sensing nutrient changes and signaling to other tissues (Arrese and Soulages, 2010; Bland, 2022; Ghosh *et al*., 2020; Huang *et al*., 2022; Kuhnlein, 2011; Martinez *et al*., 2020; Meschi and Delanoue, 2021; Sriskanthadevan-Pirahas *et al*., 2022; Stefana *et al*., 2017; Yamada *et al*., 2018). Our work demonstrates that upd3 signaling promotes hypoxia tolerance through regulation of Nos in these target tissues. Nitric oxide functions as a second messenger through activation of cGMP and PKG signaling. Previous studies have shown that NO/cGMP/PKG signaling mediates many of the physiological and behavioral adaptive responses to hypoxia in *Drosophila* embryos and larvae, including modulating gene expression, tracheogenesis, and the cell cycle (DiGregorio *et al*., 2001; Dijkers and O’Farrell, 2009; Teodoro and O’Farrell, 2003; Wingrove and O’Farrell, 1999). Nitric oxide can also mediate post-translational protein modifications through nitrosylation. Interestingly, a recent study in the *Drosophila* hematopoietic system showed that nitrosylation can enhance the unfolded protein response (Cho et al., 2024), which is a known conserved pathway induced by hypoxia (Wouters and Koritzinsky, 2008). Further studies are needed to explore whether these or other roles of nitric oxide might be important in mediating the hypoxia tolerance effects that we describe here.

Our findings support a model in which systemic responses to hypoxia, like other environmental stresses such as starvation and infection, rely on tissue-specific stress sensors and tissue-to-tissue coordination. In *Drosophila*, previous examples include sima activation in the larval fat body triggering fat-to-brain signals that suppress insulin-like peptide production under oxygen-limited conditions (Texada *et al*., 2019), and sima induction in specialized neurons modifying neuroendocrine signaling to stimulate blood cell production (Cho *et al*., 2018). Similar principles operate in mammals, where specialized tissues detect oxygen levels and trigger systemic responses—for example, oxygen sensing in keratinocytes of the epidermal layer of the skin through HIF-1α and mitochondrial metabolism triggers EPO production from the kidney to promote red blood cell production in the bone marrow (Boutin et al., 2008; Hamanaka et al., 2016), while carotid body sensing triggers neural and endocrine responses that modify respiration and circulation (Stupnikov and Cardoso, 2017). The physiology and anatomy of the fly intestine make it an ideal oxygen sensor. It is extensively innervated by a dense network of trachea that provide oxygen to support its functions and maintain its structure (Blackie et al., 2024; Li et al., 2013; Perochon et al., 2021; Tamamouna et al., 2021). This extensive tracheal network might also enable the gut to sense changes in environmental oxygen and coordinate whole-body responses via its central role as an endocrine organ. Our findings expand the view of the gut as a multipurpose environmental sensor that can detect diverse stimuli including pathogens, nutrients, and oxygen availability to coordinate systemic adaptations.

We observed a striking sexual dimorphism in this cytokine hypoxia response—although both sexes show upregulated upd3/JAK/STAT signaling in hypoxia, females require upd3 for hypoxia tolerance while males do not. Notably, sex-specific effects of upd cytokines have been reported in other contexts, including tumor growth and intestinal metabolism (Hudry et al., 2019; Wang et al., 2024), suggesting that sexual dimorphism in upd signaling may be a broader phenomenon. The effects we observe are independent of mating status, as both virgin and mated females require upd3. Studies have established that the sex determination pathway in flies can drive male-female differences in many aspects of physiology and metabolism, with whole-body metabolic differences often conferred by the sexual identity of specific cells and tissues that control and establish organismal metabolic state (Belmonte et al., 2019; Hudry *et al*., 2019; Hudry *et al*., 2016; Mank and Rideout, 2021; Millington et al., 2021a; Millington et al., 2021b; Pomatto et al., 2017; Regan et al., 2016; Regan et al., 2022; Rideout et al., 2015; Sawala and Gould, 2017; Wat et al., 2020; Wat et al., 2021). Moreover, these male-female differences in physiology confer sex dimorphic sensitivity to environmental stresses such as oxidative stress and starvation (Pomatto *et al*., 2017; Wat *et al*., 2020). Understanding how sex determination pathways might regulate upd3 signaling would be an interesting avenue to explore.

An intriguing aspect of our findings is the unexpected role of HIF-1α in restraining cytokine signaling during hypoxia. Mammalian HIF-1α is typically pro-inflammatory through control of interleukin production in mammalian immune cells (Nizet and Johnson, 2009; Palazon *et al*., 2014; Palsson-McDermott *et al*., 2015; Tannahill *et al*., 2013). However, our results suggest a more nuanced model where sima/HIF-1α can serve dual roles: activating essential hypoxia-response genes while simultaneously preventing excessive cytokine responses that could prove deleterious. The concept of restraining cytokine signaling is well-established in immunology, where unchecked cytokine production can lead to cytokine storm and tissue damage. Our work suggests that HIF-1α may function as part of this regulatory network, ensuring that hypoxia-induced cytokine responses remain within beneficial ranges. The tissue-specific nature of this regulation—with fat body sima controlling intestinal cytokine production—indicates that sima operates within a distributed regulatory system that coordinates appropriate stress responses across different organs. The molecular mechanisms underlying this inter-tissue communication remain to be determined but may involve sima-dependent regulation of secreted factors that signal from fat body to gut. Interestingly, sima has previously been shown to antagonize innate immune signaling through other pathways (Bandarra et al., 2015), suggesting that restraining inflammatory responses may be a broader function of HIF-1α depending on the specific stress conditions. Our work also points to what might be an important feature of upd biology: upd3 levels must be carefully balanced to support hypoxia tolerance. Our results show that too little upd3 compromises female survival, while excessive levels are equally detrimental. Extensive work has shown that upds can be beneficial or harmful depending on context, and our work suggests that the strength of the signaling response may be a critical factor in determining whether outcomes are adaptive or pathological.

One open question is how hypoxia controls upd3 expression in the gut—both the direct induction by hypoxia and the non-autonomous modulation by fat body sima. We found that the induction is independent of reactive oxygen species (ROS), a well-established pathway for upd3 induction under stress conditions. The induction is also independent of gut sima, contrasting with mammalian systems where HIF-1α increases similar cytokines like IL-6. The upd3 locus can integrate an array of different signaling inputs to induce gene expression in response to enteric pathogen infection (Houtz *et al*., 2017), and the hypoxia response may rely on one or more of these pathways. Metabolic changes in enterocytes could also be important. For example, dietary methionine has been shown to control upd expression through modulation of S-adenylmethionine metabolism (Obata *et al*., 2018). Interestingly, recent work has shown that in larvae, disruption of glycolysis leads to defects in larval growth and development mediated through upregulation of upd3 (Rai *et al*., 2025). Given sima’s critical role in maintaining glycolysis, the elevated gut upd3 we observe when blocking sima in the fat body may arise due to disrupted glycolysis and potentially metabolic signaling from fat body to gut.

Our findings may have important implications for understanding systemic hypoxia responses in mammals. The upd3 homolog in mammals is IL-6, a classic stress-induced inflammatory cytokine (Qing et al., 2020). Heightened IL-6 responses are observed in many disorders associated with systemic hypoxia in humans—such as sleep apnea, COPD, and COVID-19—and are thought to mediate inflammatory responses and contribute to pathology (Ayres, 2020; de Lima et al., 2016; Nadeem et al., 2013). However, our work raises the possibility that IL-6 could also be beneficial in mediating adaptive responses under physiological hypoxia. Just as we find that upd3 is essential for adaptive hypoxia tolerance in flies, IL-6 may serve important adaptive and homeostatic functions in mammals under conditions of systemic hypoxia.

## METHODS

### Fly Husbandry

*Drosophila* stocks were maintained at 25°C or 18°C. All stocks were maintained on media comprised of 100g *Drosophila* Type II agar, 1200g cornmeal, 490mg Torula yeast, 450g sugar, 1240g D-glucose, 160mL acid mixture (propionic and phosphoric acid) in 20L water. Genetic crosses were established by mating virgin females with males. All crosses and progeny were maintained at 25°C. See Table S1 for complete list of lines used.

### Hypoxia Exposure

For all hypoxia experiments, *Drosophila* were placed into an airtight chamber with a constant flow of 1% oxygen (1% oxygen/99% nitrogen) at room temperature. The flow rate was controlled using an Aalborg model P gas flow meter.

### Adult Hypoxia Survival

The duration of hypoxia exposure was tailored to each experiment to produce 50-80% survival in control animals. Males show reduced hypoxia tolerance compared to females. Accordingly, females were exposed for 24-28 hours and males for 16-18 hours (in groups of 15-25 per vial) depending on the specific experiment. The number of flies that recovered from the hypoxia exposure period were then counted.

### RU486 Treatment

Mated females were placed in vials (15-25 per vial) containing standard *Drosophila* media supplemented with 100µM RU486 in ethanol or equal volume of ethanol alone as vehicle control for 5-7 days.

### Starvation

7-day-old adult flies were placed in vials (25 per vial) containing either 0.4% agar/PBS (starved) or standard *Drosophila* media (control).

### Tissue Dissection and Microscopy

Adult female flies were surface-sterilized by rinsing with 75% ethanol in a Petri dish. Flies were then transferred to a watch glass containing ice-cold 1×PBS for dissection under a light microscope, where specific adult tissues were isolated using fine forceps. Dissected tissues were fixed in 4% paraformaldehyde (Electron Microscopy Sciences) in 1×PBS for 20 minutes at room temperature. Tissues were then stained with Hoechst 33342 dye (1:1000 dilution) for 10 minutes, followed by three 15-minute washes in 1×PBS. For mounting, tissues were placed on glass slides with coverslips using Vectashield mounting medium (Vector Laboratories Inc.). Slides were visualized using a Zeiss LSM 880 laser confocal microscope with 10× and 20× objectives and Axiovision software

### qRT-PCR

For whole-body samples, flies were transferred into 1.5mL microtubes in groups of 5 before snap freezing on dry ice. For tissue samples, groups of 5-10 female tissues were dissected in ice-cold 1xPBS and tissues immediately transferred to 1.5mL microtubes containing 500mL TRIzol reagent (Invitrogen) before snap freezing on dry ice. Total RNA was extracted from either whole flies, whole larvae, or isolated adult tissues according to manufacturer’s instructions (Invitrogen; 15596-018). RNA samples were DNase treated following manufacturer’s instructions (Ambion; 2238 G). Reverse transcription was achieved using SuperScript III (Invitrogen; 18080044). The generated cDNA was used as a template for qRT-PCR reactions (QuantStudio 6 RT-PCR system, Applied Biosystems) using the primer pairs defined in Table S2 and SYBR Green reagents. qRT-PCR amplification protocol followed a protocol for standard curve, standard ramp speed, and SYBR Green reagents. Data analysis for qRT-PCR was completed using the comparative C_T_ method (2^-ΔΔCT^). PCR data were normalized to reference genes whose expression levels were unaffected by the experimental conditions.

### Statistics

For all experiments, error bars represent Standard Error of the Mean (SEM), and p-values are the result of unpaired *t*-test or two-way Analysis of Variance (ANOVA) followed by Student’s *t*-test using GraphPad Prism (v.9). p<0.05 was considered significantly significant and indicated by an asterisk (*).

## ACKKNOWLEDGEMENTS

We thank Bruce Edgar, Erika Bach, and Bruno Lemaitre for the gift of fly stocks. Stocks obtained from the VDRC, the NIG-Fly Stock Centre, Kyoto, Japan, and the Bloomington Drosophila Stock Center (NIH P40OD018537) were used in this study. This work was supported by CIHR Project Grants (PJT-173517, PJT-195892) and NSERC DG grants to S.S.G.

---

**Figure S1:**
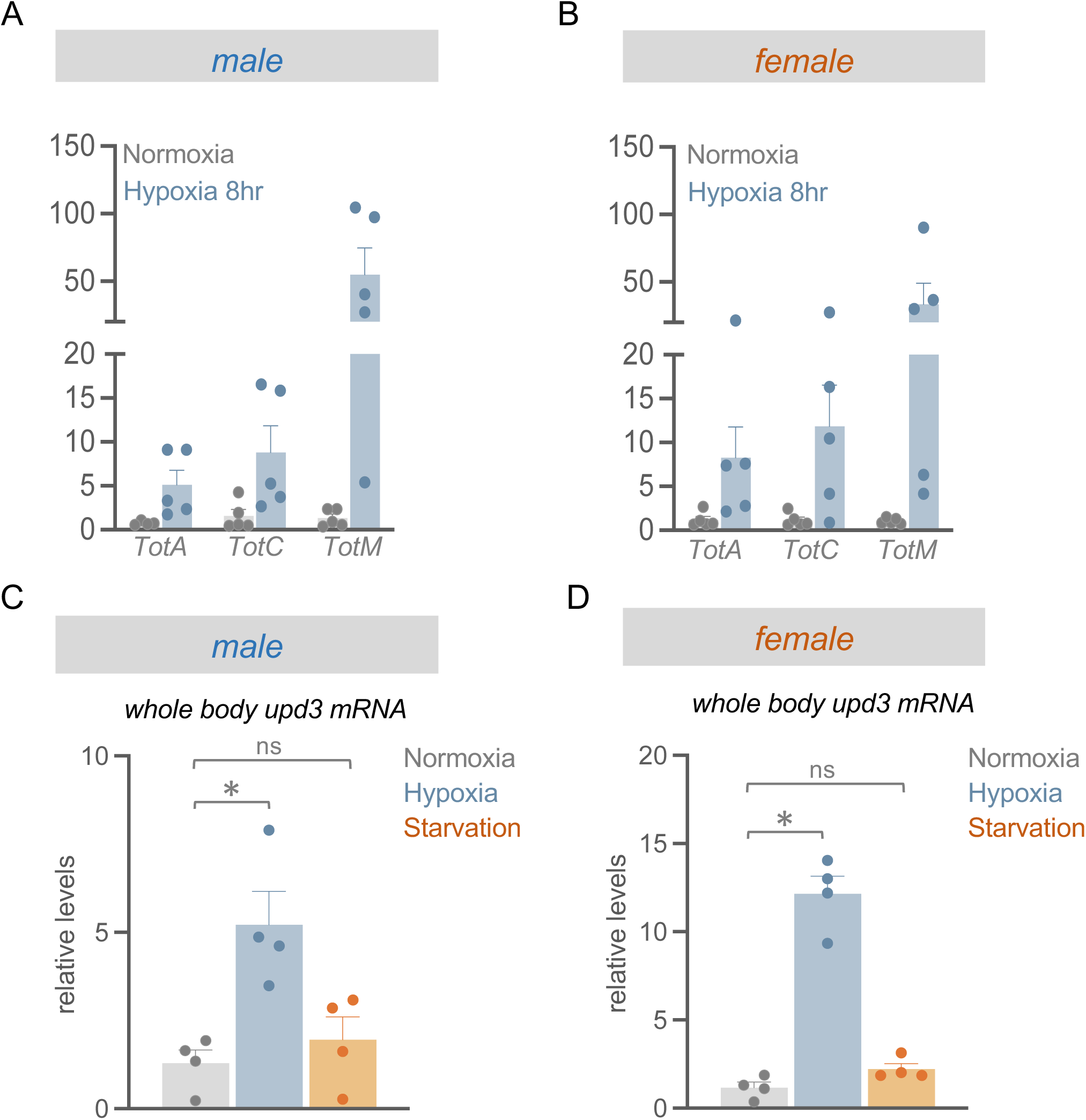
Starvation does not induce upd3 expression. A-B) *w^1118^* adult male and female flies were maintained in normoxia or exposed to 8 hours of hypoxia, then STAT-dependent *Turandot* genes were profiled. *TotA, TotC and TotM* were induced in both A) male and B) female adults. Data represents mean + SEM, N=4 independent groups of samples (5 animals per group) per experimental condition. Data points represent independent samples. C-D) *w^1118^* adult male and female flies were maintained in normoxia, exposed to 16 hours of hypoxia or 16 hours starvation. U*pd3* expression in whole animals was measured by qRT-PCR. Upon starvation, there is no significant change in *upd3* mRNA levels in either C) male or D) female adults. Data represents mean + SEM, N=4 independent groups of samples (5 animals per group) per experimental condition. *p < 0.05, ns not significant, unpaired *t*-test. Data points represent independent samples.

**Figure S2:**
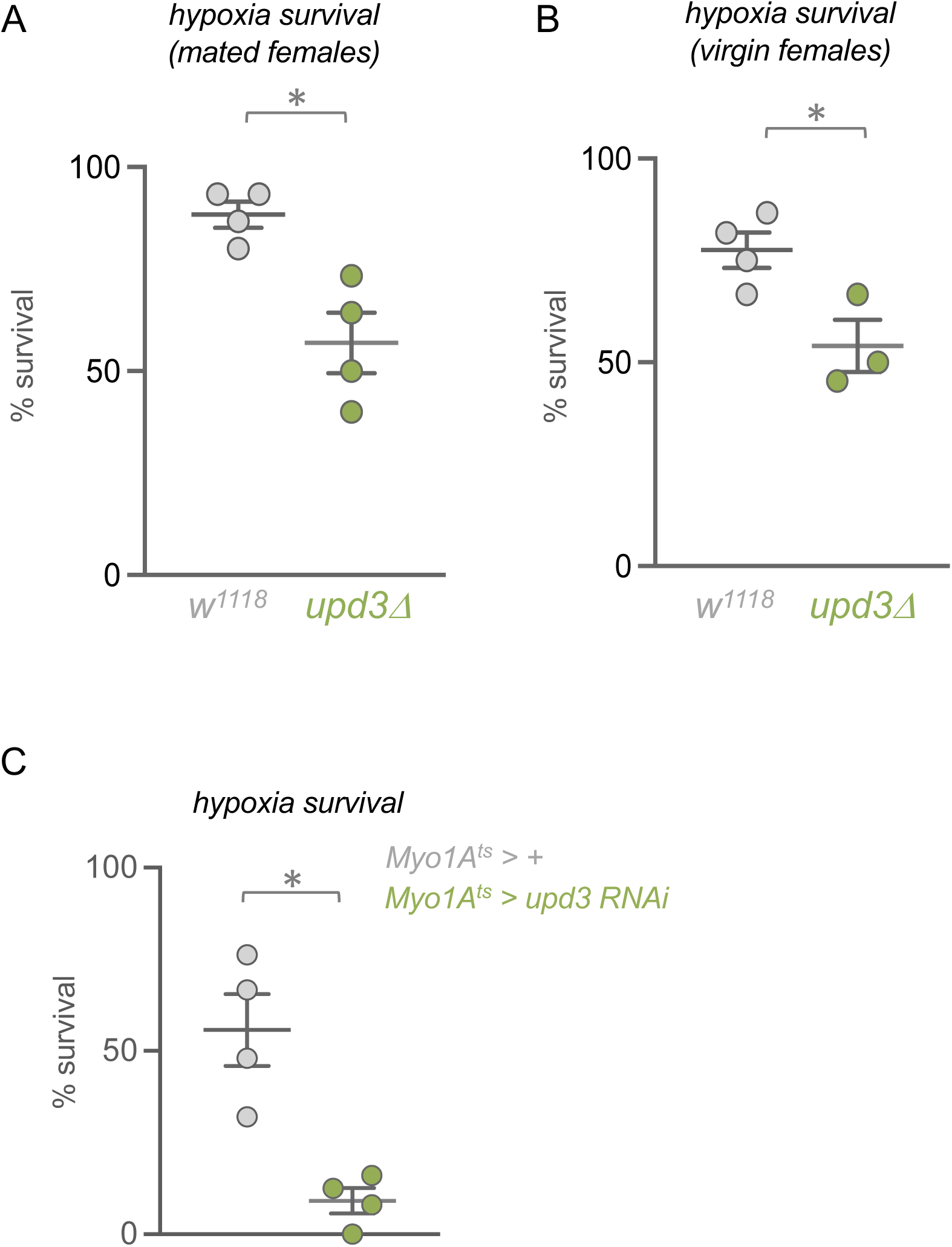
Female gut upd3 knockdown reduces hypoxia survival. Age-matched mated and virgin control (*w^1118^*) and upd3-null (*upd3Δ*) female flies were exposed to hypoxia and survival was measured. The requirement for *upd3* in hypoxia survival was not dependent on mating status in female flies – upd3-null (*upd3Δ*) A) mated and B) virgin females both showed reduced survival compared to mated or virgin controls (*w^1118^*). C) Hypoxia survival of control (*Myo1A^ts^ > +*) vs gut *upd3* RNAi (*Myo1A^ts^ > upd3-RNAi*) adult female flies. Data represents mean ± SEM, N=4 independent groups of animals (15-25 per group) per experimental condition. *p < 0.05, unpaired *t*-test. Data points represent individual samples.

**Figure S3:**
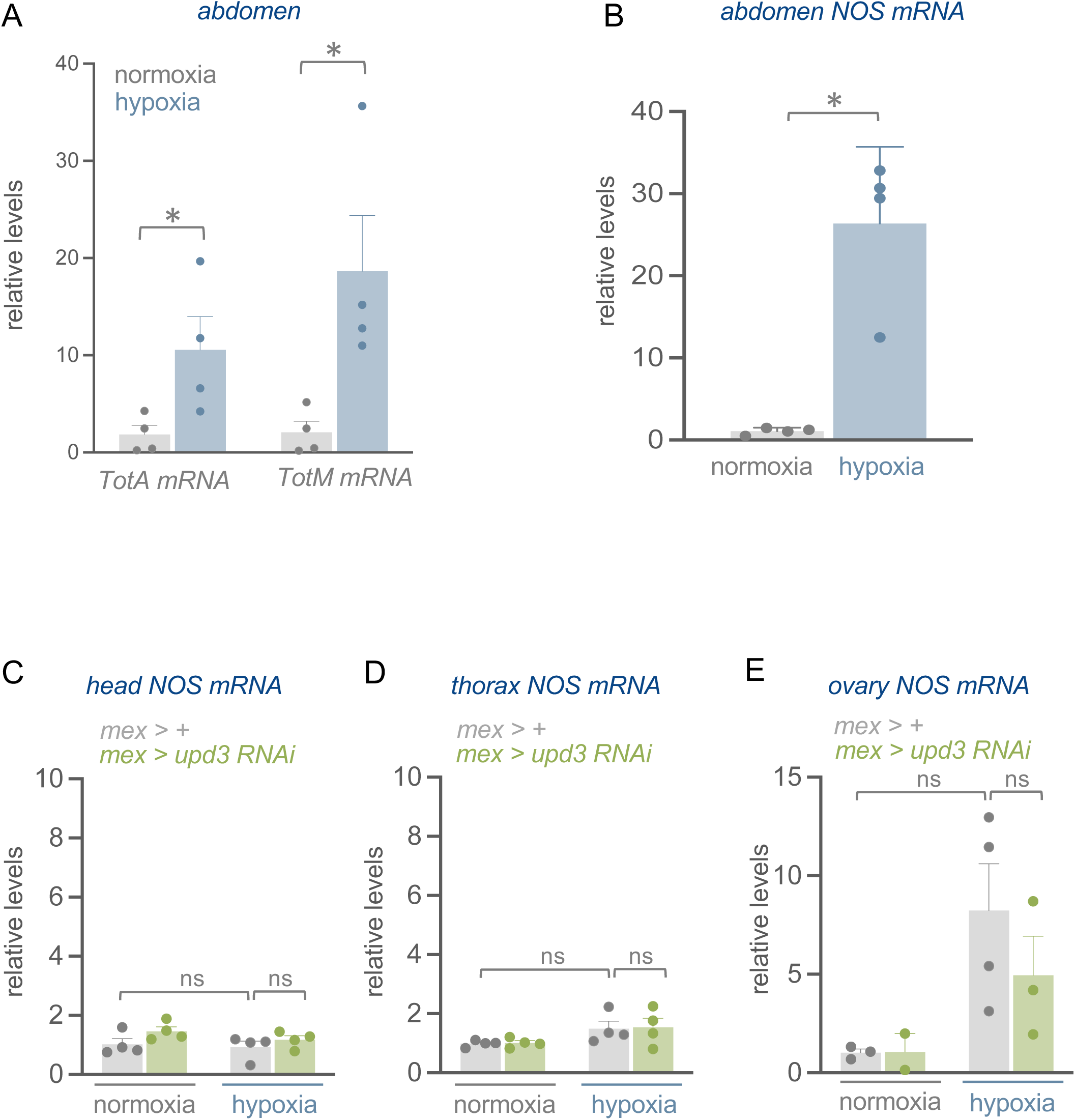
NOS regulation in hypoxia. A, B) Female flies (*w^1118^*) were exposed to 16 hours of hypoxia and abdomens isolated for gene expression analysis. A) *Turandot* genes *TotA* and *TotM* and B) *NOS* mRNA are induced in hypoxia. C-E) Intestinal upd3 knockdown (*mex > upd3-RNAi*) and control (*mex > +*) female flies were maintained in normoxia or exposed to 16h hypoxia, then tissues were collected for *NOS* mRNA gene expression analysis. Intestinal upd3 knockdown did not significantly affect *NOS* mRNA in the C) heads, D) thorax or E) ovaries in hypoxia. Data represents mean + SEM, N=2-4 independent groups of samples (5-10 tissues per group) per experimental condition. *p < 0.05, ns not significant, Student’s *t*-test following 2-way ANOVA. Data points represent independent samples.

**Figure S4:**
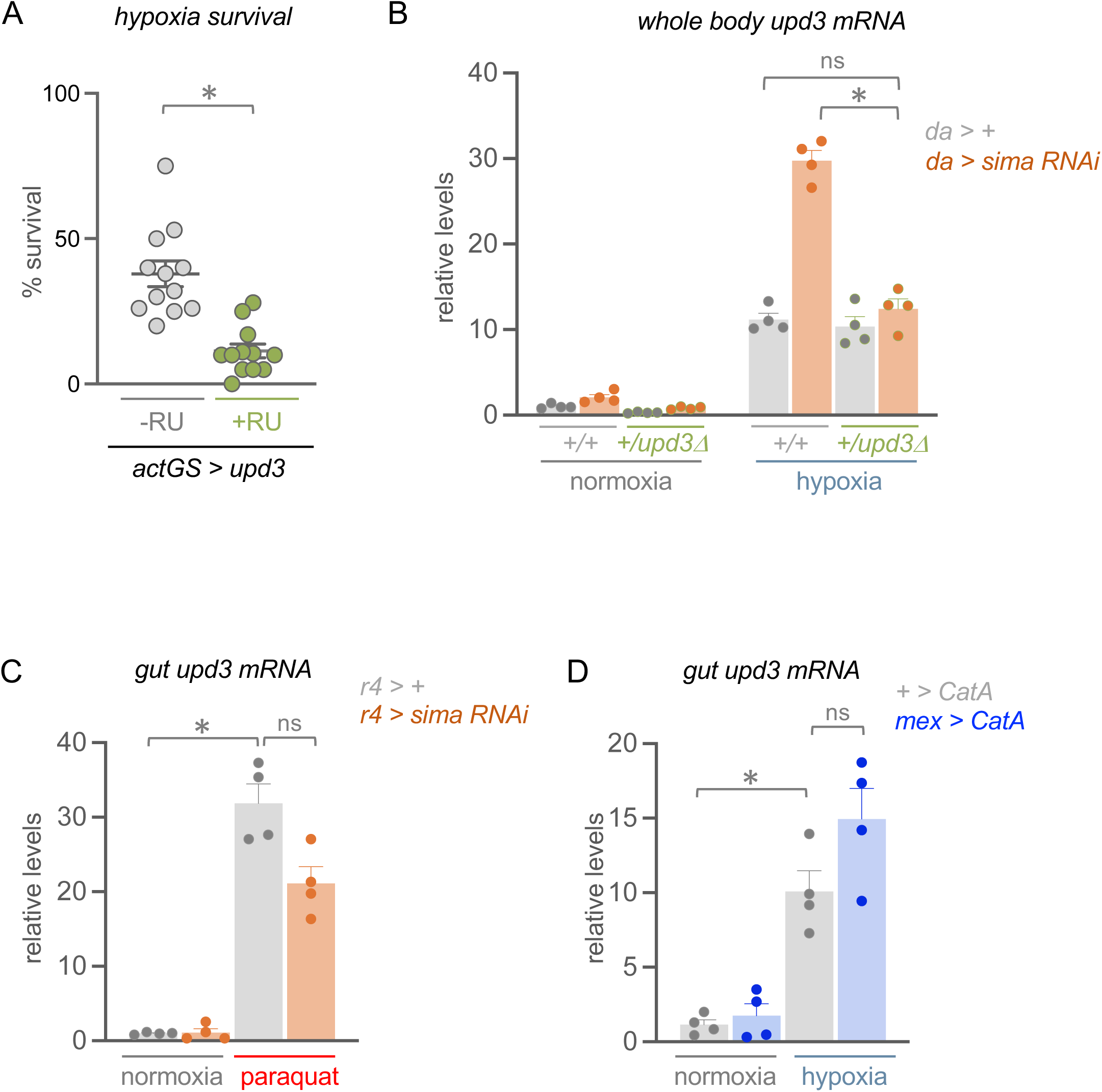
sima-dependent control of upd3. A) Hypoxia survival of control (*act-GS > upd3*, -RU486) vs *upd3* OE (*act-GS > upd3*, +RU486) adult flies. Data represents mean ± SEM, N=12 independent groups of animals (15-25 per group) per experimental condition. *p < 0.05, unpaired *t*-test. Data points represent individual samples. B) Control *(da > +)*, *upd3Δ* heterozygote (*da > upd3Δ/+*), *sima* knockdown (*da>sima-RNAi*) and *upd3Δ* heterozygote *sima* knockdown (*da > upd3Δ/+;sima-RNAi*) adult females were maintained in normoxia or exposed to 16h hypoxia. A) Hyper-induction of *upd3* mRNA levels seen in *sima* knockdown animals is abrogated in an upd3 heterozygote (*upd3Δ/+*) background. Data represents mean + SEM, N=4 independent groups of samples (5 animals per group) per experimental condition. *p < 0.05, ns not significant, Student’s *t*-test following 2-way ANOVA. Data points represent independent samples. C) Fat body sima knockdown (*r4 > sima-RNAi*) did not affect intestinal *upd3* levels compared to controls (*r4 > +*) when animals were fed oxidative stress inducing chemical paraquat. Data represents mean + SEM, N=4 independent groups of samples (5 intestines per group) per experimental condition. *p < 0.05, ns not significant, Student’s *t*-test following 2-way ANOVA. Data points represent independent samples. D) Knockdown of ROS scavenger Catalase A (*mex > CatA*) failed to block intestinal induction of *upd3* compared to controls (*+ > CatA*) in hypoxia. Data represents mean + SEM, N=4 independent groups of samples (5 intestines per group) per experimental condition. *p < 0.05, ns not significant, Student’s *t*-test following 2-way ANOVA. Data points represent independent samples.

**Table S1.** *Drosophila* stocks used in this work.

| Name | Genotype | Source |
| --- | --- | --- |
| upd3Gal4,UAS-GFP | +; upd3-gal4, UAS-GFP/Cyo; + | Gifted from Erica Bach |
| upd3Δ | w, upd3Δ; +; + | BDSC (55728) |
| UAS-upd3 | w<sup>1118</sup>; UAS-upd3; + | Gifted from Bruno Lemaitre |
| UAS-upd3 RNAi | +; +; UAS-upd3 RNAi | VDRC (27136/GD) |
| UAS-sima RNAi | +; UAS-sima RNAi; + | VDRC (106187/KK) |
| UAS-NOS RNAi | yv; UAS-NOS RNAi; + | BDSC (57700) |
| UAS-STAT92E RNAi | +; UAS-STAT92E RNAi; + | VDRC (43866/GD) |
| UAS-CatA | w; UAS-CatA; + | BDSC (24621) |
| mexGal4<sup>ts</sup> | mex-GAL4/CyO; tubgal80ts, UAS-nlsGFP/TM6B | Gifted from Bruce Edgar |
| daGal4 | w<sup>1118</sup>; +; da-gal4 | BDSC (8641) |
| r4Gal4 | w<sup>1118</sup>; +; r4-gal4 | BDSC (33832) |
| desatGal4 | w; desatGal4/Cyo; + | BDSC (65404) |
| mexGal4 | w<sup>1118</sup>; mex-gal4; + | BDSC (91368) |
| Myo1AGal4 | w; Myo1A-gal4; + | Gifted from Bruce Edgar |
| Myo1AGal4<sup>ts</sup> | w; Myo1AGal4 tubGal80ts UAS-GFP | Jiang et al., 2009 |
| daGS | w<sup>1118</sup>; da-gal4 GeneSwitch; + | - |
| actGS | w<sup>1118</sup>; Act-gal4 GeneSwitch; + | - |

**Table S2.**
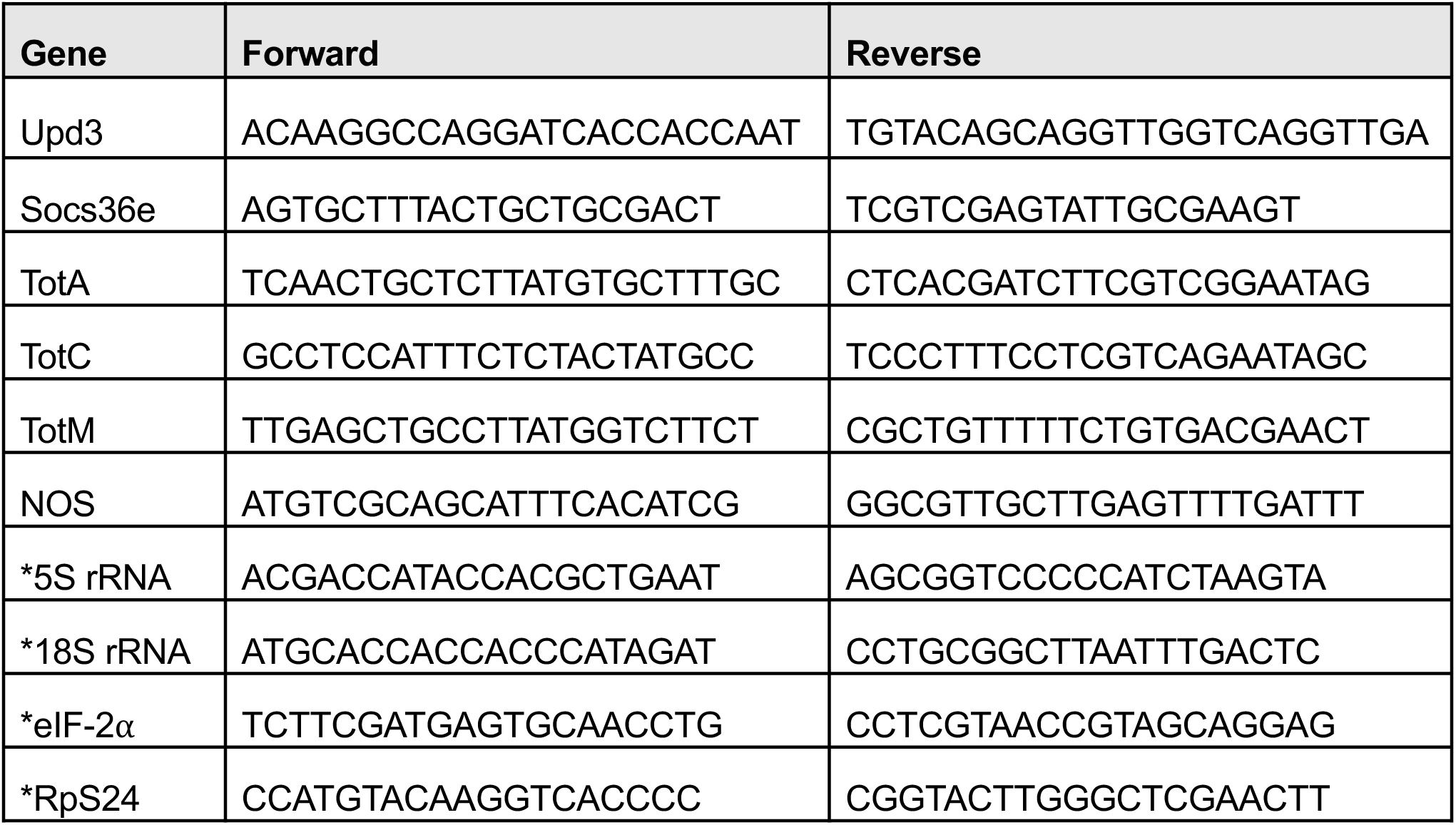
Primers used for qRT-PCR. All primers listed in 5’-3’ direction. Reference genes used in this study are indicated by an asterisk (*).

## Notes

### Competing Interest Statement

The authors have declared no competing interest.

